# Disruption of metazoan gene regulatory networks in cancer alters the balance of co-expression between genes of unicellular and multicellular origins

**DOI:** 10.1101/2023.04.20.537744

**Authors:** Anna S Trigos, Felicia Bongiovanni, Magnus Zethoven, Richard Tothill, Richard Pearson, Tony Papenfuss, David L Goode

## Abstract

Metazoans inherited genes from unicellular ancestors that perform essential biological processes such as cell division, metabolism and protein translation. Functioning multicellularity requires careful control and coordination of these unicellular genes to maintain tissue integrity and homeostasis. Gene regulatory networks (GRNs) formed during metazoan evolution to regulate conserved biological processes are frequently altered in cancer, resulting in over-expression of unicellular genes. We propose an imbalance in co-expression of unicellular (UC) and multicellular (MC) genes is a driving force in cancer. To investigate, we combined gene co-expression analysis to infer changes to GRNs in cancer with protein sequence conservation data to distinguish genes with UC and MC origins. Co-expression networks created using RNA sequencing data from 31 tumour types and normal tissue samples were divided into modules enriched for UC genes, MC genes or a mix of both (Mixed UC-MC modules). The greatest differences between tumour and normal tissue co-expression networks occurred within Mixed UC-MC modules. In particular, MC and UC genes not commonly co-expressed in normal tissues formed distinct co-expression modules seen only in tumours. The degree of rewiring of genes within Mixed UC-MC modules increased with both tumour grade and stage. Mixed UC-MC modules were enriched for somatic mutations in cancer genes, particularly copy-number amplifications, suggesting an important driver of the rewiring observed in tumours are copy-number changes. Overall, our study shows the greatest changes to gene co-expression patterns during tumour progression occur between genes of MC and UC origins, implicating the breakdown and rewiring of metazoan gene regulatory networks in cancer development and progression.

**Author summary:** Multicellular organism cells follow certain rules that control and coordinate their growth and behavior. This happens because gene regulatory networks formed during the evolution of multicellularity to control the activity of genes inherited from unicellular ancestors. Cancer cells disobey these rules, growing and dividing in a competitive fashion analogous to that of colonial unicellular organisms. Here, we test the hypothesis that breakdown of gene regulatory networks enforcing multicellularity drives cancer progression by investigating 31 tumour types. Based on sequence similarity, genes were categorized as having origins in either unicellular or multicellular species. We found that the balance of expression unicellular and multicellular genes changes dramatically in cancer. Genes expressed together in normal tissues stop being co-expressed in tumors, while unicellular and multicellular genes that would not normally be expressed together in normal tissues become highly co-expressed. This phenomenon is more pronounced in cancers at more advanced stages, and sometimes occurs in association with gain or loss of parts of certain chromosomes. Our work indicates disruption and rewiring of gene regulatory networks that evolved to enforce multicellularity drives cancer progression by upsetting the carefully coordinated balance in the activity of unicellular and multicellular genes.

## Introduction

Changes to transcriptional regulation played a major role in the evolution of complex multicellularity^1, 2^. Early metazoan species inherited genes from their unicellular ancestors that underpinned core biological functions such as cell division, motility and metabolism. Stable multicellularity required fine control over where and when such processes occurred. Expanded gene regulatory networks (GRNs) evolved to coordinate activity of unicellular genes and sustain the cooperative growth crucial for functional multicellularity^3, 4^. The essentiality of unicellular genes for cellular homeostasis meant a simple on/off mode of regulation would not suffice. Metazoans need to maintain expression of unicellular genes at safe levels, and periodically upregulate them for development, wound healing, stress responses and other processes. Metazoan GRNs evolved to balance expression of genes with unicellular and multicellular origins, leading to their consistent co-expression across healthy tissues^5, 6^.

Many hallmarks of cancer involve unravelling of core features of multicellularity^7^, including reduced cell-cell adhesion, lack of coordinated cell division, altered metabolism and breakdown of tissue differentiation. The atavism hypothesis of cancer proposes reactivation of conserved transcriptional programs inherited from ancestral unicellular species enhances the adaptability and replicative potential of tumour cells, linking changes in gene regulation to loss of multicellularity in cancer^8–11^.

Increasing molecular evidence supports a role for changes to the balance of expression between unicellular and multicellular genes during the formation and progression of cancer. In earlier work, we found pervasive down-regulation of genes of multicellular origin and selective upregulation of genes of unicellular origin across solid cancers^5, 12^, since independently confirmed^13^. Similar approaches showed both genetic and transcriptional alterations accumulate in genes involved in multicellularity during selection for accelerated growth and metastatic potential in breast cancer xenografts^14^, in highly conserved genes in myeloma upon exposure to chemotherapy^15^ and in recently-evolved paralogs of cell cycle control genes in cancer cell lines^16^. An example of the functional impacts is the stress-induced activation of error prone, stress-activated DNA repair mechanisms conserved with bacteria in a range of tumours, facilitating drug resistance^17, 18^ and leaving detectable mutational signatures^19, 20^.

Human protein-protein interaction (PPI) and gene regulatory networks show layering of genes by evolutionary age, with highly conserved genes of unicellular origin found closer to the centre and more recently evolved genes on the peripheries^21, 22^. This creates structural vulnerabilities that can lead to cancer^23^. We found transcriptional regulators forming hubs connecting genes with unicellular origins with genes of multicellular origins in human GRNs are frequently enriched for somatic mutations in cancer^24^, implicating the dysfunction of such hubs as important drivers of tumour formation and progression. Intriguingly, somatic amino acid substitutions in known cancer driver genes sometimes match the dominant or fixed allele in unicellular eukaryotes^25–27^ causing direct reversion to ancestral protein sequences that may alter oncogenic pathways.

Here, we investigate how disruption of metazoan GRNs leads to imbalance in expression of unicellular and multicellular genes in cancer. First, protein sequence orthology data were used to assign evolutionary age categories to human genes. Genes conserved in multiple single-celled species were placed in the unicellular category. Otherwise, genes were labeled as being multicellular if orthologs could only be found in other metazoan species. We performed gene co-expression analysis to infer gene regulatory networks for 31 types of solid cancers. Overlaying evolutionary gene age categories onto co-expression networks found major changes in co-expression of unicellular and multicellular genes occurs during tumour formation and progression. Unicellular and multicellular genes not usually co-expressed in normal tissues come together to form distinct co-expression modules in tumours that often include known cancer driver genes. The activity and degree of rewiring within these modules increases with both tumour grade and degree of malignancy.

Altogether, disruption and rewiring of metazoan GRNs is a common and important aspect of tumour progression that causes imbalance in the co-expression of unicellular and multicellular genes. Our framework demonstrates a new way to combine sequence conservation and co-expression analysis to study the links between the evolution and multicellularity and cancer.

## Results

### Tracking changes in co-expression between unicellular and multicellular genes in cancer

Alterations to the structure of metazoan GRNs that evolved to support multicellularity should manifest as differences in patterns of co-expression of unicellular and multicellular genes in tumours relative to normal somatic tissues. To investigate, we constructed a three-step pipeline to combine to sequence orthology data with gene co-expression networks to study how genes of unicellular or multicellular origin are co-regulated in cancer (Fig 1A).

**Figure 1:**
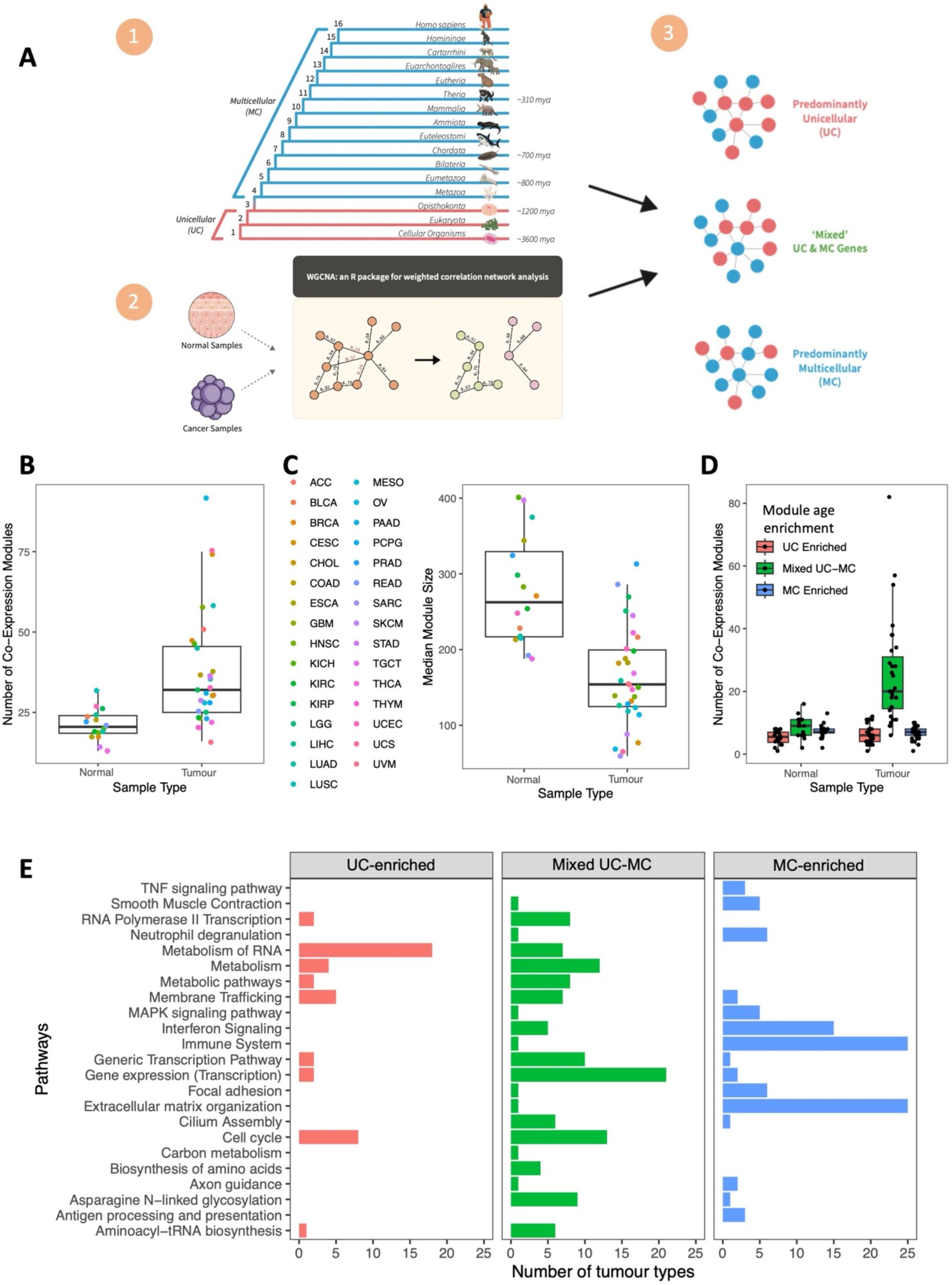
WGCNA-derived gene co-expression modules in tumour and normal samples. **(A)** Identification and classification of gene co-expression modules. (1) Phylostratigraphy was applied to assign unicellular or multicellular origins to genes. (2) WGCNA was used to cluster genes into co-expression modules based on correlation in expression. (3) Modules were classified according to the degree of enrichment of genes of unicellular or multicellular origin. **(B)** Summary of number of modules per TCGA cohort, with points coloured according to tumour type. **(C)** Mean number of genes in tumour and normal modules per TCGA cohort. Colour key is the same as in (B). **(D)** Distribution of numbers of Unicellular-enriched, Multicellular-enriched and Mixed UC-MC modules across TCGA cohorts. **(E)** KEGG and Reactome terms found to be most frequently enriched in modules across all TCGA cohorts. The total numbers of tumour types containing a Unicellular, Mixed UC-MC or Multicellular module enriched for each term are shown on the x-axis.

We first assigned evolutionary ages to human genes using phylostratigraphy, which employs phylogenetics and multiple sequence alignments to estimate the point in evolutionary time where a gene emerged^6^. Phylostratigraphic analysis revealed cancer genes tend to be highly conserved, and enriched in particular for genes that emerged in unicellular organisms and during early stages of metazoan evolution^28^. We divided species in 16 clades representing the major stages of evolution from single-celled organism to modern humans (Fig 1A, panel 1), as we have previously done^5^. Briefly, for each human gene, protein sequence alignments from OrthoMCL^29^ were used to identify the most distant clade from human in which species with high-confidence orthologs could be found (Methods). Genes with orthologs in bacteria or single-cell eukaryotes (clades 1-3) were designated as having unicellular origins (UC genes). Genes with orthologs in other multicellular species only were classified as being of multicellular origin (MC genes). Clade assignments are provided in Supplementary Data.

Next, we inferred the structure and topology of GRNs in both tumours and normal somatic tissues by applying the weighted gene co-expression network analysis (WGCNA) algorithm^30^. WGCNA identifies ‘modules’ of genes with correlated expression across multiple samples (Fig 1A, panel 2), implicating underlying coordinated regulatory mechanisms exist that connect those genes into identifiable co-expression modules. WGCNA co-expression modules were generated for each of the 31 solid tumour types with available RNA-sequencing data in The Cancer Genome Atlas (TCGA). For the 16 TCGA cohorts with data for at least 10 normal samples, we generated both tumour and normal gene co-expression modules (bladder urothelial carcinoma - BLCA, breast invasive carcinoma - BRCA, colon adenocarcinoma - COAD, oesophageal carcinoma - ESCA, head and neck squamous cell carcinoma - HNSC, kidney chromophobe - KICH, kidney renal clear cell carcinoma - KIRC, kidney renal papillary cell carcinoma - KIRP, liver hepatocellular carcinoma - LIHC, lung adenocarcinoma - LUAD, lung squamous cell carcinoma - LUSC, prostate adenocarcinoma - PRAD, rectum adenocarcinoma - READ, stomach adenocarcinoma - STAD, thyroid carcinoma - THCA, uterine corpus endometrial carcinoma – UCEC). Here, gene expression data from matched pairs of tumour-normal samples were used to identify co-expression modules. For cohorts with no normal samples available, only tumour co-expression modules were built, using gene expression data from all available tumour samples.

Finally, co-expression modules were classified according to prevalence of unicellular or multicellular genes (Fig 1A; Methods) into three categories. Modules enriched in UC genes were defined as being Unicellular-enriched, while modules enriched in MC genes were defined as Multicellular-enriched. Modules not enriched in either UC or MC genes were placed into the Mixed UC-MC category and represent regions of human GRNs where there is communication between genes of different evolutionary ages.

In total, 16 to 92 modules were obtained per TCGA tumour-type cohort, for a total of 1,503 modules (337 from normal tissues and 1166 from tumour tissues). On average there were more modules in tumour samples (37.61 per cohort) than in normal samples (21.06 per cohort) (Fig 1B) (two-sided Wilcoxon test p-value = 4.77×10^-5^). This trend held even when only cohorts with matched tumour and normal samples were considered (two-sided Wilcoxon test p-= 6.87×10^-5^) (Supp Fig S1). The number of modules detected per cohort was not a function of sample size (Supp Fig S2). Tumour modules were typically about half the size of normal modules, with an average of 163.4 genes per tumour module versus an average of 278.1 genes per normal module (two-sided Wilcoxon test p-value = 1.63×10^-5^) (Fig 1C). Again, this was true when comparing only cohorts with matched normal samples (two-sided Wilcoxon test p-value = 6.87×10^-5^) (Supp Fig S3) and was independent of cohort size (Supp Fig S2). Generally, the modules obtained from WGCNA provide comprehensive coverage of the transcriptome from TCGA cohorts, with more than 90% of genes assigned to modules in 28 of 31 tumour types (Supp Fig S4). These patterns are consistent with the greater heterogeneity seen between tumours and breakdown of links between different GRN regions in cancer.

The higher numbers of co-expression modules derived from tumour samples was due to an excess of modules belonging to the Mixed UC-MC category, which contain a balanced mix of unicellular and multicellular genes. An average of 24.97 Mixed UC-MC modules were obtained per TCGA tumour cohort, ranging from 6-82 per tumour type (Fig 1D and Supp Table S1). Collectively, 774 of the 1166 (66.38%) modules found across all tumour cohorts combined were Mixed UC-MC. In marked contrast, only 137/337 (40.65%) of the combined modules from all normal cohorts together were Mixed UC-MC (Supp Fig S5) (Fisher Exact test p=4.87×10^-17^, Odds Ratio=2.88 (2.23-3.73)). Unicellular-enriched modules comprised 185/1166 (15.87%) of modules from tumor samples whereas 207/1166 (17.75%) of tumour modules were Multicellular-enriched (Supp Table S1). In contrast, 84/337 (24.93%) modules from normal cohorts were Unicellular-enriched while 116/337 (34.42%) were Multicellular-enriched, significantly different proportions from tumour modules (Fisher Exact Test p =1.46×10^-16^) Most tumour cohorts had similar numbers of Unicellular-enriched (mean of 5.97) and Multicellular-enriched (mean of 6.68) modules, comparable to normal cohorts, which averaged 5.25 Unicellular-enriched and 7.25 Multicellular-enriched modules, respectively. (Fig 1D and Supp Table S1).

To understand their functional roles all co-expression modules were tested for enrichment of terms from the Kyoto Encyclopedia of Genes and Genomes^31^ and Reactome^32^ using gProfiler^33^. Unicellular-enriched modules were associated with basic cellular processes such as translation, metabolism and cell cycle, consistent with the highly conserved nature of the genes in those modules. Modules from the Multicellular-enriched category were largely associated with functions specific to multicellularity, such as immunity and the extracellular matrix (Fig 1E). Interestingly, Mixed UC-MC modules were enriched for terms related to transcription and mRNA processing as well as core processes such as metabolism and cell cycle, consistent with this class of modules representing processes used by metazoans regulate gene expression of unicellular pathways.

Gene co-expression analysis suggests GRNs in cancers can be divided into regions predominantly composed of genes from unicellular or multicellular ancestors, or regions connecting both types of genes. The differences in the prevalence and size of Mixed UC-MC modules between tumour and normal cohorts suggests the onset of cancer is accompanied by substantial changes in co-expression of unicellular and multicellular genes.

### Gene co-expression analysis implicates substantial rewiring of regulatory links between unicellular and multicellular genes in cancer

To assess the frequency and magnitude of rewiring between UC and MC genes into new co-expression modules, we devised a metric to measure the extent to which modules in tumors contain unique assemblages of genes relative to their matched normal counterparts. Our metric, which we refer to as ‘Novelty’ score, is designed to capture the relative degree of change in co-expression between conditions represented by a given module. Briefly, it reflects the number of genes within a given tumour module that were also found together in a module from the corresponding normal tissue type (Fig 2A; Methods). Based on this score, tumour modules were designated as low, medium or high Novelty based on their overlap with normal modules from the corresponding tissue type. Low Novelty modules preserve correlation in expression between tumour and normal tissues; for example, transcriptional programs that retain activity in tumours. Medium or high Novelty modules have fewer co-occurring pairs with normal modules and thus better capture co-expression patterns present in tumours but not in the corresponding normal tissue.

**Figure 2:**
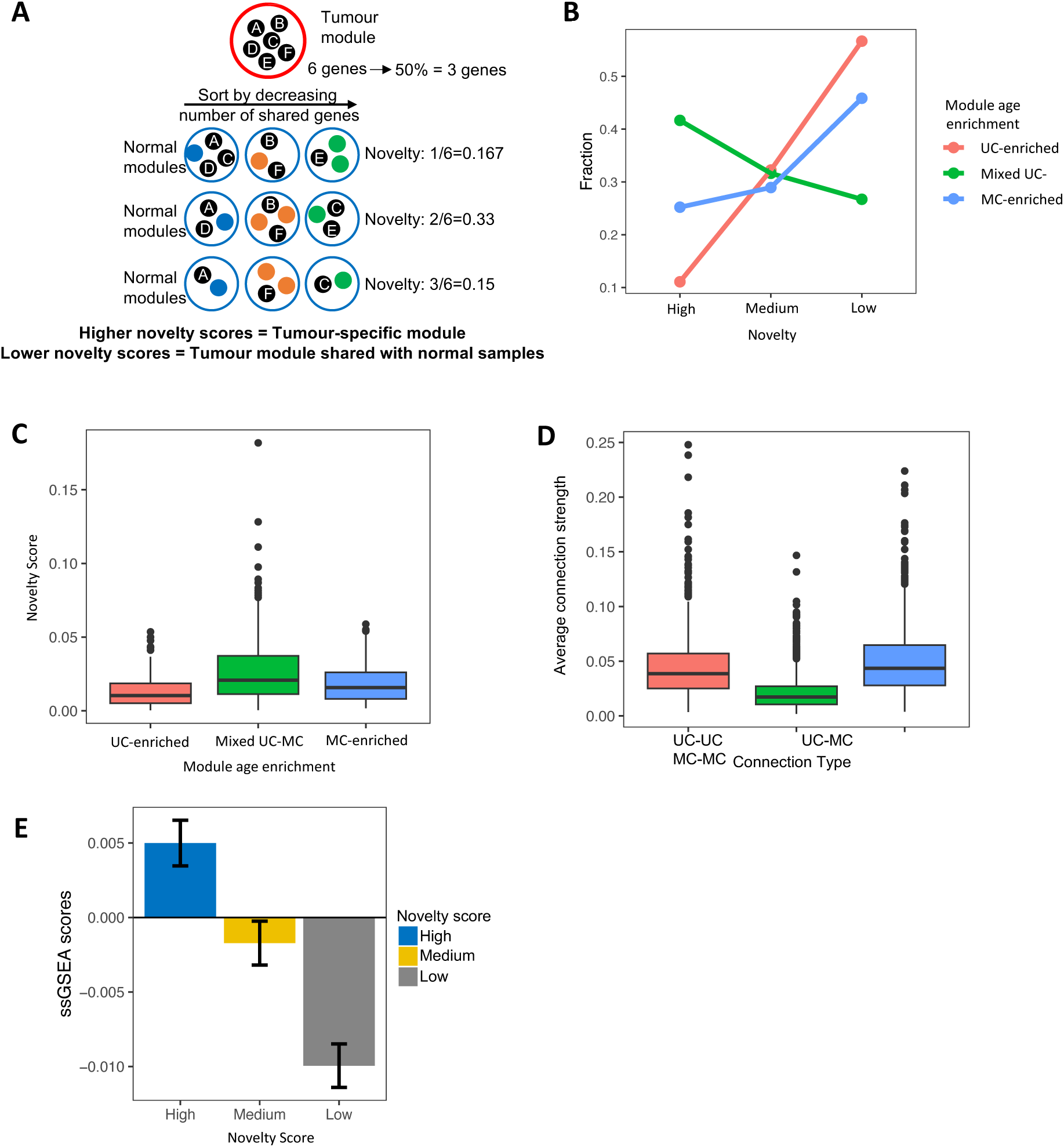
Measuring divergence in co-expression between WGCNA modules from tumours and normal tissues. **(A)** Demonstration of calculation of module novelty scores based on overlap in gene content between tumour and normal co-expression modules. **(B)** Fraction of UC-enriched (pink), Mixed UC-MC (green) and MC-enriched (blue) modules with High, Moderate or Low novelty, combined across all 16 TCGA cohorts with matching normal tissue data. **(C)** Distribution of novelty scores for tumour modules enriched in unicellular genes (UC-enriched; pink), containing a mix of unicellular and multicellular genes (Mixed UC-MC; green) or enriched in multicellular genes (MC-enriched; blue). Data are combined across all 16 TCGA cohorts with matching normal tissue data. **(D)** Average edge weights for edges between pairs of Unicellular genes (UC-UC; pink), between Unicellular and Multicellular genes (UC-MC; green) and between pairs of Multicellular genes (MC-MC; blue), for all tumour modules from all TCGA cohorts combined. **(E)** Single-sample GSEA scores for tumour modules with High (blue), Medium (yellow) and Low (grey) novelty scores, across all 16 TCGA cohorts with matching normal tissue data.

Novelty of modules was tightly associated with module age. Overall, 56.66% of all UC-enriched modules and 45.85% of MC-enriched tumour modules across all cohorts were low novelty (Fig 2B), indicating correlations in expression between pairs of genes that are both from unicellular phylostrata or both from multicellular phylostrata are more likely to be preserved in cancer. Mixed UC-MC tumour modules had higher novelty scores than modules enriched for UC or MC genes (one-sided Wilcoxon test p-values = 1.22×10^-11^ and 4.80×10^-5^, respectively) (Fig 2C, Supp Fig S6). Averaging across cohorts, 41.66% of Mixed UC-MC modules were high novelty (Fig 2B), forming the majority of the high novelty tumour modules identified in tumour cohorts (Supp Fig S7).

Mixed UC-MC modules appear to capture new gene co-expression patterns not present in normal tissue, indicating regulatory links between UC and MC genes are disrupted and rewired during tumor development. We hypothesized if there is rewiring of gene regulatory networks during tumor development, there would be substantial heterogeneity in co-expression of UC and MC genes between patients. In agreement, edge weights assigned by WGCNA between UC and MC genes (UC-MC) were weaker than edges between unicellular genes (UC-UC) or between multicellular genes (MC-MC) when we averaged the strength of these connection types in each module (one-sided Wilcoxon p-values < 2.2×10^-16^ in both cases), indicating higher levels of heterogeneity and/or transcriptional plasticity in co-expression of unicellular and multicellular genes in cancer (Fig 2D). Modules in all gene age enrichment categories contain both unicellular and multicellular genes, meaning while connections between unicellular and multicellular genes are enriched in Mixed UC-MC modules, they also contribute significantly to UC-enriched and MC-enriched modules (Supp Fig S8).

Finally, measuring the combined expression of the genes in each module in tumours using single-sample Gene Set Enrichment Analysis (ssGSEA)^34^ revealed high Novelty modules had the highest ssGSEA scores, followed by medium then low Novelty modules (Fig 2E). On average, in the upper quartile of ssGSEA scores in each tumor type, 31.53% of modules were high novelty modules, 27.03% of medium novelty modules, compared to just 20.79% of low novelty modules (Supp Fig S9). This shows high and medium novelty modules result from active gene expression, as opposed to peripheral contamination from noise in the data.

Overall, these findings indicate the greatest differences in co-expression between tumours and normal samples occur between genes found in Mixed Unicellular-Multicellular modules. Mixed UC-MC modules display both high levels of expression in tumours and major differences in co-expression patterns from normal tissues, as evidenced by their higher Novelty scores. This fits a model where the balance in co-expression between unicellular and multicellular genes is consistently altered in tumours through changes in the underlying gene regulatory networks.

### The presence of somatic mutations impacts the composition and topology of tumour gene co-expression modules

Cancer progression is defined by accumulation of somatic mutations that enhance fitness, often by causing dramatic changes to regulation of transcription. Our prior work showed somatic mutations to key hubs in GRNs disrupt communication between unicellular and multicellular genes^24^. To examine how somatic mutations affect co-expression of unicellular and multicellular genes, we overlaid data on copy-number alterations, coding insertions and deletions (indels) and predicted deleterious point mutations from TCGA onto modules generated by WGCNA.

First, we calculated the percentage of known cancer drivers from the COSMIC Cancer Census gene list^35^ in each tumour co-expression module. We matched genes with the tumour type in which they are considered to be drivers. Novelty score was associated with fraction of known driver genes, with high Novelty modules containing the highest percentage of COSMIC Cancer Census genes (Fig 3A) compared to medium (one-sided Wilcoxon test p-value=7.01×10^-7^) and low novelty modules (p=9.49×10^-12^). Furthermore, there was overall strong correlation between module Novelty score and percentage of known cancer drivers (Supp Fig S10) (Spearman rho: 0.72; p-value = 4.52×10^-23^). Gene age enrichment category was also associated with presence of known cancer genes, as Mixed UC-MC modules had a higher percentage of COSMIC Cancer Census genes than UC-Enriched or MC-Enriched modules (one-sided p-values=0.11 and 0.032, respectively) (Fig 3B). These findings are consistent with somatic driver mutations leading to formation of novel gene expression associations between unicellular and multicellular genes.

**Fig 3:**
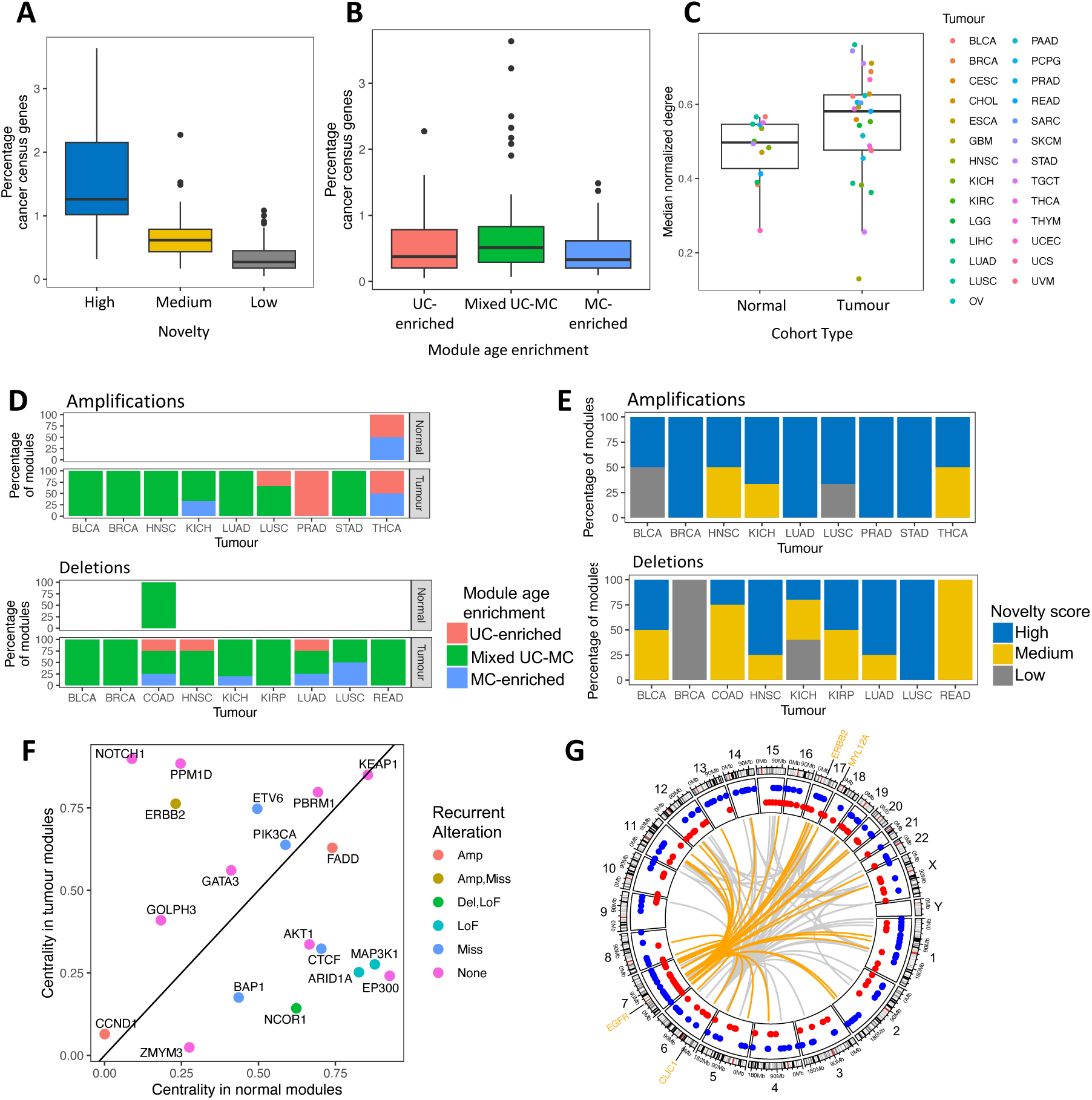
The locations and frequencies of driver genes within tumour gene co-expression modules. **(A)** Percentages of COSMIC Cancer Census gene list that are found in High (blue), Medium (yellow) or Low (grey) novelty modules, across tumour modules with matched normal samples. **(B)** Percentage of COSMIC Cancer Census genes found in UC-enriched (pink), Mixed UC-MC (green) or MC-enriched (blue) modules, across tumour modules with matched normal samples. **(C)** Normalized degree of recurrently amplified genes within WGCNA co-expression modules from normal (left) and tumour (right) cohorts, coloured according to tumour type. **<u>(D)</u>** Gene age enrichment distribution for modules where >10% of genes are recurrently amplified (top) or recurrently deleted (bottom) within the indicated tumour types. Bars show overall percentages of such modules that are UC-enriched, Mixed UC-MC or MC-enriched. **(E)** Novelty score distribution for modules where >10% of genes are recurrently amplified (top) or recurrently deleted (bottom) within the indicated tumour types. **(<u>F</u>)** Degree centrality in normal modules versus tumour modules for known driver genes in breast cancer. Points coloured by the type of recurrent alteration observed in that gene in the BRCA TCGA cohort. **(G)** CIRCOS plot of the ‘purple ‘module from the TCGA low-grade glioma cohort. Points show genomic locations of genes with unicellular (red) or multicellular (blue) origins. Gray lines indicate *cis* and *trans* connections within and between chromosome for the 100 most strongly co-expressed gene pairs in this module. Locations of the most highly connected genes (EGFR, CLIC1, ERBB2 and MYL12A) are indicated, with the strongest 50 connections to the EGFR oncogene highlighted in orange.

Given the importance of copy-number aberrations (CNAs) in driving gene expression changes in cancer we assessed the prevalence of recurrently amplified or deleted genes, i.e., those gained/lost in >10% of patients within a given subtype (Methods). Tumour modules harbouring recurrently amplified genes tended to be more central in the network (i.e. have stronger edge weights) overall than did modules without any recurrently mutated genes (one-sided Wilcoxon test p-value = 0.024) (Fig 3C) suggesting amplifications enhance the strength and consistency across patients of co-expression between unicellular and multicellular genes. In contrast deletions did not have consistent effects on gene centrality in tumor co-expression modules (Supp Fig S11).

The enrichment of recurrently mutated genes in co-expression modules from tumours was primarily due to their associations with amplifications and deletions. Of the 337 modules derived from normal tissues, in only 2 (0.59%) were at least 10% of genes recurrently amplified in cancer. In contrast, 51 of 1166 (4.37%) tumour modules contained >=10% genes recurrently amplified in the corresponding cancer type (one-sided Fisher Exact p=1.56×10^-4^, Odds Ratio=0.13 (0.00-0.43)). Likewise, only 1 normal module was enriched in genes affected by deletions in cancer while across all tumour types 39/1166 (3.34%) of co-expression modules contained >10% recurrently deleted genes (one-sided Fisher Exact p=4.32×10^-4^, Odds Ratio=0.086 (0.00-0.43)). These enrichments are consistent with the capacity for copy-number alterations (CNAs) to dramatically elevate or reduce expression of genes in cancer^36^.

The greatest prevalence of recurrent copy-number alterations was seen in high Novelty modules from the Mixed UC-MC category. In 7 of 9 tumour types with modules with more than 10% of genes harboring recurrent amplifications, the majority of these modules were of the Mixed UC-MC category. Similarly, of the modules associated with deletions the majority were Mixed UC-MC in 8 of the 9 cohorts with modules with more than 10% of genes harboring recurrent deletions (Fig 3D). Amplifications were also more prevalent in modules with high Novelty scores. Across all 9 tumour cohorts, high novelty modules comprised 50% or more of the modules containing recurrently amplified genes (Fig 3E). In contrast, modules with recurrently deleted genes had a broader mix of Novelty scores (Fig 3E). Therefore, some amplifications appear to cause altered co-expression of unicellular and multicellular genes, leading to formation of high Novelty Mixed UC-MC modules.

We assessed how the impact of recurrently mutated genes on the transcriptomes in tumours may be mediated by their topological positions within co-expression networks represented by our WGCNA modules. We hypothesized genes driving formation of distinct co-expression modules in tumours migrate to a more central and interconnected positions compared to their original location in normal modules. Conversely, genes involved in the fragmentation of normal modules during tumourigenesis would become less connected and migrate to the periphery of tumour modules. To test this, we calculated the centrality of each gene after normalizing edge weights across modules (Methods), revealing structural differences between tumor and normal modules.

Marked changes in centrality of genes between normal and tumour co-expression modules in key driver genes from the breast cancer (BRCA) cohort of TCGA. Multiple known breast cancer driver genes had large changes in centrality (Fig 3F) that aligned with their established roles in tumourigenesis. Oncogenes (e.g. NOTCH1 and ERBB2) tended to increase in centrality in tumour modules while tumour suppressors (e.g. BAP1, AKT1 and EP300) decreased in centrality. Similar findings emerged in other solid tumour types (Supp Fig S12).

Across all cohorts amplifications had the strongest changes in centrality. Recurrently amplified genes increased centrality in 20 of the 27 TCGA cohorts where they could be found (Supp Fig S13) suggesting that upon amplification they drive form key hubs in distinctive new co-expression modules in tumours. Deletions were also tied to changes in gene centrality, but the impact of gene deletions appears to be context dependent as the effects on centrality vary by cohort (Supp Fig S14). The centrality of recurrently deleted genes was on average reduced from normal tissues to tumours in 15 of the 29 TCGA cohorts examined but increased instead in 10 cohorts. It is important to note here WGCNA builds modules around both positive and negative correlations in expression between genes^30^. The loss of strong regulators can result in strong negative correlations captured by co-expression patterns specific to certain tumour types.

Gain or loss of large chromosomal segments can lead to simultaneous but coincidental expression changes to multiple linked genes in addition to the selected driver gene or genes^37^. While that may explain some associations between recurrently amplified genes and co-expression modules in tumours, we observed many cases where recurrent amplifications were linked to co-expression changes in involving loci across the genome. An example is the ‘purple’ module from the low-grade glioma (LGG) cohort of TCGA, which harbours the EGFR oncogene a well-known driver of disease progression in glioma^38^ (Figure 3G). Higher activity of this high Novelty Mixed UC-MC module was associated with poor survival in the LGG cohort (Supp Fig S15). The LGG purple module contained genes from every chromosome except chrY and strong co-expression of EGFR with genes across the genome (Fig 3G). The most highly connected gene in this module is CLIC1, a chloride ion channel with high expression and potential therapeutic and prognostic value in adult glioma^39, 40^. Our work links CLIC1 expression with EGFR activity in LGG.

### The extent of altered co-expression of unicellular and multicellular genes increases during tumour progression

Widespread changes in co-expression patterns between unicellular and multicellular genes in tumours indicates this phenomenon is a general feature of solid cancers. Rewiring of links connecting unicellular and multicellular in GRNs could result in the emergence of many features that accompany tumorigenesis^11, 23^. If so, such changes to co-expression of unicellular and multicellular genes would be expected to become more pronounced as tumours progress to more malignant states.

To investigate, we tracked the differences in gene co-expression between distinct stages of tumour progression, by sorting samples according to tumour grade and degree of malignancy (Methods). Comparison were performed in three solid tumour RNA-sequencing data sets: (1) data from TCGA prostate cancer (PRAD) cohort, and sets of (2) benign skin nevi and melanoma samples^41^ as well (3) malignant pheochromocytomas and benign pheochromocytomas^42^. Gene co-expression modules were generated using WGCNA for each category of samples within each data set. To assess how gene co-expression changed during tumour progression we used our Novelty score metric to compare modules from high-grade or malignant samples to those from low-grade or benign samples from the same tumour type. Where available, module Novelty scores were also computed for benign and low-grade tumours against matched normal tissues, to understand how co-expression changed during early stages of tumour development. Comparison of gene age enrichments in modules was performed to determine whether the greatest changes in co-expression occurred in UC-enriched, Mixed UC-MC or MC-enriched modules.

For the TCGA PRAD cohort, Novelty scores were calculated for modules from high-grade primary tumours (annotated as Grade groups 4 and 5) compared to the modules derived from low-grade primary tumours (Grade groups 1-3), as well for low-grade tumour modules against modules derived from adjacent non-diseased prostate gland tissue (Methods). The greatest increase in novelty scores occurred in high-grade primary prostate tumours relative to low-grade tumours (Fig 4A). In contrast, the novelty scores for modules from low-grade tumours were relatively modest compared to modules from normal tissue samples. Overall, just 23.08% of low-grade modules had medium or high novelty but 75.96% of high-grade modules had medium or high novelty (Fig 4B), indicating the majority of changes to co-expression occur during later stages of development of primary tumours in prostate cancer. In both comparisons, Mixed UC-MC modules had the highest Novelty scores, demonstrating the greatest rewiring in co-expression during prostate cancer progression happens between unicellular and multicellular genes (Fig 4A). High-grade modules were predominantly Mixed UC-MC across all novelty categories (21.31%, 36.61% and 37.70% high grade modules were of low, medium and high novelty modules and mixed UC-MC) were Mixed UC-MC across all novelty categories (Fig 4C).

**Figure 4:**
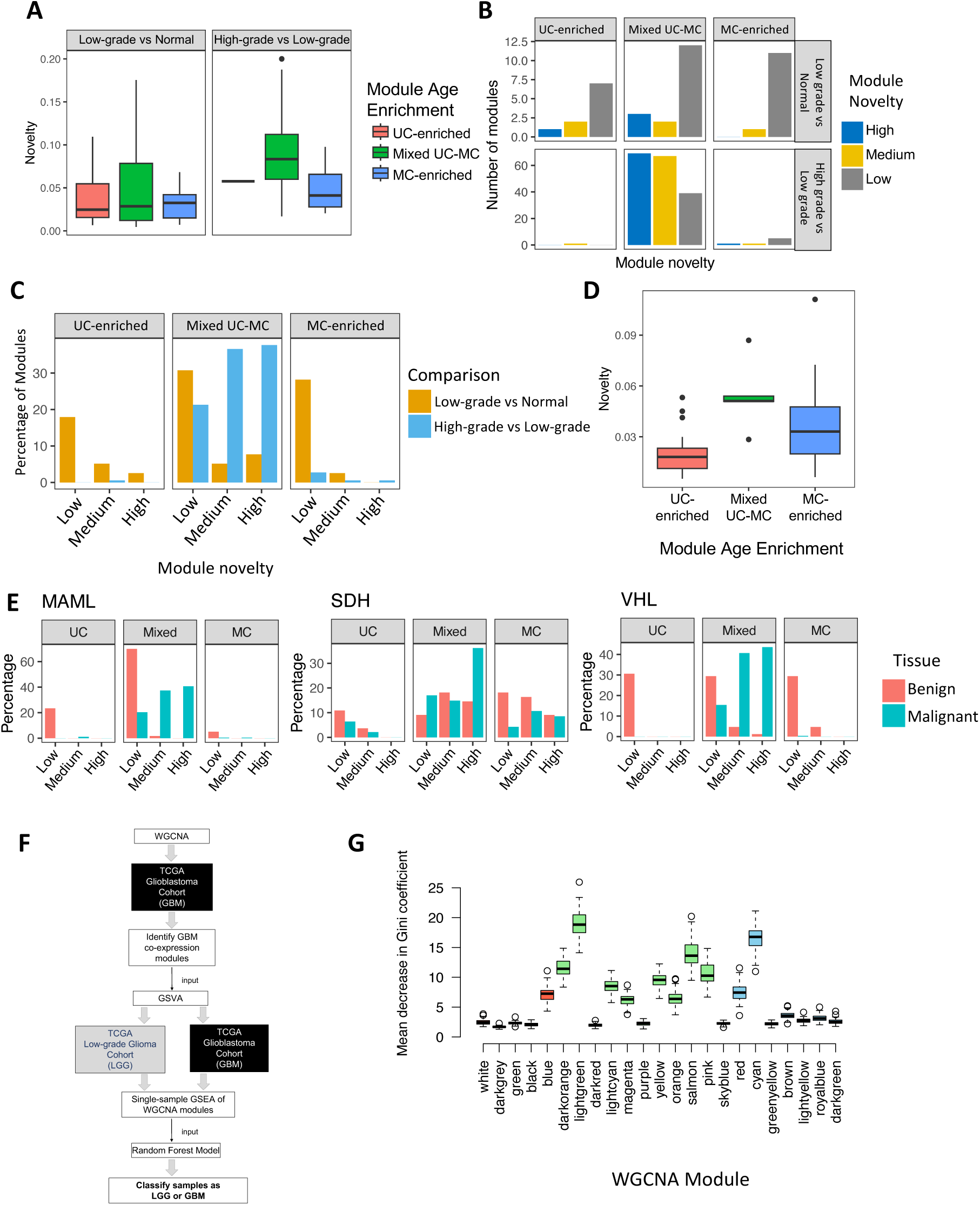
Changes in co-expression between unicellular and multicellular genes during tumour progression. **<u>(A)</u>** Novelty scores for Unicellular-enriched (pink), Mixed Unicellular-Multicellular (green) and Multicellular-enriched (blue) co-expression modules derived from low-grade prostate tumours (Gleason Grade groups 1 and 2) as compared to co-expression modules from normal (non-diseased) prostate tissue (left) and for co-expression modules from high-grade tumours (Gleason Grade groups 4 and 5) as compared to co-expression modules from low-grade prostate tumours (right). Data from TGCA PRAD cohort. **(B)** Number of UC-enriched, Mixed UC-MC and MC-enriched modules with low, medium or high novelty scores from low-grade (top) and high-grade (bottom) prostate tumours. **(C)** Percentage of co-expression modules with low, medium or high novelty scores from comparison low-grade prostate tumours to normal prostate tissues (gold) and from comparison of high-grade to low-grade prostate tumours (blue), stratified by gene age enrichment category (UC-enriched, Mixed UC-MC or MC-enriched). **(D)** Novelty score distribution for UC-enriched (pink), Mixed UC-MC (green) or MC-enriched (blue) co-expression modules derived from melanoma as compared to co-expression modules derived from benign nevi, using data from Badal et al. **(E)** Percentage of co-expression modules with low, medium or high novelty scores for benign tumours compared to normal adrenal gland tissue (pink) and malignant tumours compared to benign tumours (blue), stratified by gene age enrichment category and pheochromocytoma subtype. **(F)** Schematic of process of using Random Forest model to classify adult glioma samples as LGG or GBM based on ssGSEA score of WGCNA modules derived from the TCGA glioblastoma cohort. **<u>(G)</u>** Power of co-expression modules derived from the TCGA glioblastoma cohort to distinguish low-grade from glioblastoma samples, expressed as distribution of mean decrease in Gini Coefficient for 100 replicates of a Random Forest model. Gene age enrichment category is indicated by color: UC-enriched (pink), Mixed UC-MC (green) or MC-enriched (blue).

To understand how co-expression of unicellular and multicellular genes differs between benign and malignant lesions we analysed a cohort published by Badal et al^41^, which contains RNA-sequencing data from 50 primary melanomas of the skin and 27 benign skin growths known as nevi. We found Mixed UC-MC modules from melanomas had significantly higher Novelty scores when compared to modules derived from benign nevi than did Unicellular-enriched or Multicellular-enriched modules (Fig 4D) (one-sided Wilcoxon test p-values = 8.62×10-4 and 0.059, respectively).

To investigate the full spectrum of co-expression changes during progression from normal tissue to benign lesions to tumours, we created separate sets of WGCNA modules from RNA-sequencing data from benign and malignant pheochromocytoma samples^42^, as well as healthy adrenal tissue (Methods). Tumour samples were further divided into the MAML, SDH and VHL subtypes, which have distinct driver mutations. Across all subtypes, modules derived from benign tumours showed low Novelty as compared to modules from normal adrenal glands, regardless of gene age enrichment. However, novelty scores were markedly higher when comparing modules between malignant and benign pheochromocytomas with Mixed UC-MC modules on average having the higher novelty than Unicellular-enriched and Multicellular-enriched modules (Fig 4E) (Wilcoxon test one-sided p-values= 5.88×10^-8^, 5.15×10^-5^, 9.62×10^-15^, comparing Mixed UC-MC with UC-enriched in MAML, SDH and VHL samples, respectively, and 0.0019, 0.10, 1.27×10^-13^, comparing Mixed UC-MC with MC-enriched in MAML, SDH and VHL samples, respectively). The melanoma and pheochromocytoma data thus support the notion gene regulatory connections between unicellular and multicellular genes become progressively more altered as tumour cells develop into more malignant states.

### Mixed Unicellular-Multicellular gene co-expression modules best distinguish low-grade from high-grade glioma tumours

The excess of high-Novelty Mixed Unicellular-Multicellular modules in high-grade and malignant tumours indicated the degree of alteration of co-expression of unicellular and multicellular genes distinguishes tumours at different stages of progression. We next hypothesized that these co-expression modules would provide significant discriminant power to classify samples based on their degree of malignancy. To test this, we focused on the low-grade glioma (LGG) and glioblastoma (GBM) datasets of TCGA. LGGs almost inevitably progress to the high-grade and fatal GBM. LGGs and GBMs thus represent linear stages in glioma evolution with clear differences in degree of malignancy. ssGSEA scores were calculated for the 23 WGCNA modules obtained from the GBM cohort, to measure the combined level of expression of genes within each module in each of the 512 samples from the LGG cohort and the 153 samples from the GBM cohort. Module ssGSEA scores were then used to train an RF model to distinguish between LGG and GBM samples from TCGA (Fig 4F). RF classification provides a means to determine which co-expression modules are most closely associated with disease stage in glioma. One hundred independent iterations were performed using random splits of training and testing samples each time (Methods).

Median Area Under the Curve (AUC) of the Receiver Operating Characteristic (ROC) curves for the 100 RF models based on GBM module ssGSEA scores was 0.814 (Interquartile range of 0.818 – 0.894). This was comparable to the performance of RF models trained on the status of mutations normally used to grade adult glioma (median AUC=0.814), though models based on mutations had much more variable performance (Interquartile range of 0.524 – 0.879) (Supp Figure S16). Thus, changes to gene co-expression have power to accurately classify LGG from GBM samples across patients.

We expected Mixed UC-MC modules would be most distinctive between LGG and GBM and therefore carry the greatest weight in the RF classifiers. To investigate, we calculated mean decrease in the Gini coefficient for each GBM WGCNA module, a measure the importance of a given variable to an RF model. In total, 11 of the 22 GBM modules had a mean decrease in the Gini coefficient greater than 5, indicating they made a substantial contribution to the predictive power of the RF model (Fig 4G). Of these, 8 modules were Mixed UC-MC with 40-60% genes from multicellular phylostrata (72.7%; Fisher test p-value = 0.0296). These results were subsequently validated by random forests trained and tested on orthogonal RNA-sequencing data from GBM and LGG patients from the Chinese Glioma Genome Atlas (CGGA). ssGSEA scores for TCGA GBM modules could also distinguish LGG from GBM samples in the CGGA (Median AUC = 0.667, IQR 0.640 – 0.717) (Supp Figure S16). In the CGGA RF models, 6 of the 8 modules (75%) with a decrease in Gini >5 had a Mixed UC-MC composition (Supp Figure S17). All 6 of these Mixed UC-MC modules also had a decrease in Gini >5 in the TCGA RF models.

The bulk of the predictive power of our gene co-expression-based RF classifiers thus derive from the Mixed UC-MC class of WGCNA modules. This underscores the importance of altered co-expression and therefore altered regulatory links between genes of unicellular and multicellular origins in glioma as it progresses from low-grade to high-grade disease. Such changes are biologically meaningful, as distinctive patterns of gene co-expression clearly distinguish LGGs from GBMs in two independent cohorts.

## Discussion

We demonstrate pronounced alterations to co-expression of genes of unicellular and multicellular origins are pervasive across tumour transcriptomes. This brings about an imbalance in co-expression of unicellular and multicellular genes relative to normal, non-malignant tissues. There are a number of important implications for how GRNs that evolved in metazoans to enforce multicellularity are disrupted and rewired in cancer.

First, GRNs get fragmented into smaller and more isolated subnetworks in cancer. More gene co-expression modules were found in tumours but on average they contained fewer genes than co-expression modules derived from normal tissues. To quantify the degree of change represented by tumour co-expression modules we devised a new gene overlap metric termed Novelty score. The biggest Novelty scores tended to occur in modules in the Mixed Unicellular-Multicellular category. In contrast, tumour modules enriched for unicellular or multicellular genes (UC-Enriched or MC-Enriched) tended to have low Novelty scores. In the context of inferred structures of metazoan GRNs, our analyses suggest major changes occur to GRNs in cancer at the interface between unicellular and multicellular genes to disrupt communication between regions of predominantly unicellular genes or predominantly multicellular genes. This effect likely accompanies and amplifies the dysregulation caused by mutations to GRNs(elife ref).

Mixed UC-MC modules likely have important functional impacts rather than being by-products of transcriptional noise. Mixed UC-MC modules with high Novelty scores also showed a trend of higher expression in tumours and higher prevalence of known driver genes relative to UC-Enriched and MC-Enriched modules. Somatic mutations may act to force the common expression of pairs of unicellular and multicellular genes that would not normally be co-expressed in healthy somatic tissues^16, 24^. Amplifications in particular can alter expression of multiple genes simultaneously and thereby rapidly disrupt metazoan GRNs, perhaps explaining their strong enrichment in Mixed UC-MC modules. However, many Mixed UC-MC modules did not contain known driver genes, indicating environmental, epigenetic and other factors not explored here cause imbalanced co-expression of unicellular and multicellular genes in tumours as well.

Alterations to co-expression between unicellular and multicellular genes continues as cancer progresses. We observed increased dissimilarity in Mixed UC-MC modules from high-grade and malignant tumours relative to their low-grade and benign counterparts, respectively. In contrast, the composition of UC-Enriched and MC-Enriched modules was comparatively stable across tumour stages. This held across a diverse range of tumours from prostate, skin, brain and the adrenal gland. We propose the disruption and rewiring of metazoan GRNs is progressive, accelerating at later stages of tumour development to create altered patterns of co-expression of unicellular and multicellular genes that enhance malignancy. Investigation of co-regulatory links between unicellular and multicellular genes may thus uncover new prognostic biomarkers and therapeutic opportunities. The predictive value of Mixed UC-MC modules for distinguishing tumour grade in glioma using Random Forests highlights this potential.

Co-expression analysis carries an inherent degree of uncertainty related to biological and technical variation between samples. While this may impact the accuracy of individual modules, the consistent alteration to co-expression of unicellular and multicellular genes across 32 diverse types of solid tumours indicates a common and reproducible biological effect across cancers. TCGA samples contain both cancerous cells and non-malignant stroma, and some co-expression modules likely capture features of the tumour microenvironment. However, as Mixed UC-MC modules are strongly enriched for processes up-regulated in tumours and many are associated with known driver mutations, we expect they largely stem from tumour-intrinsic gene expression changes. Isolation of tumour cells through single-cell approaches may offer further clarity as techniques for inferring gene regulatory networks *de novo* from single-cell expression data improve.

The atavism hypothesis of cancer is often interpreted as representing a simple loss of regulation of unicellular genes in cancer. The reality is of course more nuanced, as many genes of multicellular origins are also activated and play key roles in cancer. We propose rather than just suppressing ancient transcriptional programs, metazoan GRNs serve to strike a balance in the co-expression of unicellular and multicellular genes. Integrating activities of core biological programs inherited from unicellular ancestors with novel functionalities and regulatory circuits that emerged with multicellularity enabled evolution of the stunning array of forms and functions seen in modern metazoans. When regulatory mechanisms enforcing this integration break down, cancer can emerge. Our findings thereby provide a more complex and flexible view of atavism, one in line with recent developments in the field towards a more stepwise and selective process underlying the loss of multicellularity in cancer.

This work creates a foundation for investigating the full implications and mechanisms of how the loss of features of multicellularity drive cancer. Integrating other ‘omics modalities such as proteomic, epigenetic and chromatin conformation data along with comparative analysis of gene co-expression in unicellular and basal metazoan species with tumours present fertile ground for future studies.

## Materials and methods

### Phylostratigraphy

We used phylostratigraphy^6^ to determine the evolutionary ages of genes, as we have previously done^5^. Briefly, human genes were mapped onto a phylogenetic tree of 16 phylostrata based on sequence similarity, spanning genes found across all organisms (Phylostratum 1) to those specific to humans (Phylostratum 16). Genes with orthologs in bacteria or single-cell eukaryotes (clades 1-3) were designated as having unicellular origins (UC genes). Genes with orthologs in other multicellular species only were classified as being of multicellular origin (MC genes).

### Gene expression datasets

Tumour and normal gene expression datasets were obtained from The Cancer Genome Atlas (TCGA) for 31 tumour types (adrenocortical carcinoma - ACC, bladder urothelial carcinoma - BLCA, breast invasive carcinoma - BRCA, cervical squamous cell carcinoma and endocervical adenocarcinoma - CESC, cholangiocarcinoma - CHOL, colon adenocarcinoma - COAD, oesophageal carcinoma - ESCA, glioblastoma multiforme - GBM, head and neck squamous cell carcinoma - HNSC, kidney chromophobe - KICH, kidney renal clear cell carcinoma - KIRC, kidney renal papillary cell carcinoma - KIRP, brain lower grade glioma - LGG, liver hepatocellular carcinoma - LIHC, lung adenocarcinoma - LUAD, lung squamous cell carcinoma - LUSC, mesothelioma - MESO, ovarian serous cystadenocarcinoma - OV, pancreatic adenocarcinoma - PAAD, pheochromocytoma and paraganglioma - PCPG, prostate adenocarcinoma - PRAD, rectum adenocarcinoma - READ, sarcoma - SARC, skin cutaneous melanoma - SKCM, stomach adenocarcinoma - STAD, testicular germ cell tumors - TGCT, thyroid carcinoma - THCA, thymoma - THYM, uterine corpus endometrial carcinoma - UCEC, uterine carcinosarcoma - UCS, uveal melanoma - UVM).

For cohorts with data for at least 10 normal samples (BLCA, BRCA, COAD, ESCA, HNSC, KICH, KIRC, KIRP, LIHC, LUAD, LUSC, PRAD, READ, STAD, THCA, UCEC), gene expression data from the available matched pairs of tumour-normal samples were used.

To avoid unreliability in the levels of expression of genes, only genes with a count per million (cpm) greater than 1 in either all tumour or all normal samples, respectively, were included.

### Building gene co-expression modules

WGCNA^30^ version 1.61 was used to build gene co-expression modules. As suggested by WGCNA, we used hierarchical clustering based on gene expression to discard dissimilar samples and ensure high consistency between the samples used for the analysis, only keeping paired samples that survived the preprocessing steps. Genes were filtered by WGCNA’s preprocessing function goodSamplesGenes().

Next, the standard procedure to generate gene co-expression modules using WGCNA was followed. Briefly, a large unsigned adjacency matrix was built based on the absolute values of the Pearson correlation coefficients between genes using the adjancency() function. To measure interconnectedness, the topological overlap between genes was calculated using the TOMsimilarity() function. The resulting weighted co-expression network generated by WGCNA represents the genes (nodes) and the strength of co-expression associations between them (edges). Next, hierarchical clustering was performed using hclust() with the ‘average’ agglomerative method and cutreeDynamic() was used to cut the resulting tree to partition the graph into modules. Modules were required to have at least 30 genes. All modules passed the quality checks performed by WGCNA. The grey module corresponds to genes not assigned to a particular module and was excluded from analyses.

This procedure to build modules were performed independently for each tumour and normal cohort. Besides the membership of genes to modules, we also extracted the interactions between genes in a module by subsetting the initial gene co-expression network built with all genes, resulting in a weighted network of all-against-all genes per module.

### Assigning ages to modules

To assign ages to modules, one-sided Fisher enrichment tests were performed for each module using the number of UC and MC genes in the module and the overall number of UC and MC genes in the gene expression cohort dataset used to generate the modules. Correction for multiple testing was performed using the Benjamini-Hochberg procedure, and significance was achieved when the adjusted p-value was less than 0.05. Modules were classified as UC if they were enriched in UC genes, MC if they were enriched in MC genes, and Mixed UC-MC if they were not enriched in either.

### Functional enrichment of modules

We used gProfiler^33^ R package to perform functional enrichment of modules using gene sets from the Kyoto Encyclopedia of Genes and Genomes^31^ and Reactome^32^ using the g:GOst function, which performs over-representation analysis. The top functional enrichment categories were selected for each module. Pathways unrelated to cancer and general cancer pathways were excluded. Specifically, we excluded the following terms: Huntington disease, Human cytomegalovirus infection, Alzheimer disease, Staphylococcus aureus infection, Central carbon metabolism in cancer, Hepatitis C, MicroRNAs in cancer, Proteoglycans in cancer, Measles, Pertussis, Small cell lung cancer, Hepatocellular carcinoma, Viral myocarditis, Parkinson disease, Insulin resistance, Human papillomavirus infection, Diseases of signal transduction, Chronic myeloid leukemia, HIV Infection, Pathways in cancer, Loss of Function of SMAD4 in Cancer, Vibrio cholerae infection, Platinum drug resistance, Hypertrophic cardiomyopathy (HCM),Gastric cancer, Non-small cell lung cancer, Toxoplasmosis, Epstein-Barr virus infection, Renal cell carcinoma, Shigellosis, Basal cell carcinoma, Colorectal cancer, Endometrial cancer, Human immunodeficiency virus 1 infection, Kaposi sarcoma-associated herpesvirus infection, Herpes simplex virus 1 infection, AGE-RAGE signaling pathway in diabetic complications, Defective CFTR causes cystic fibrosis, Dilated cardiomyopathy (DCM), Hepatitis B, Central carbon metabolism in cancer, MicroRNAs in cancer, Proteoglycans in cancer, Small cell lung cancer, Pathways in cancer, Gastric cancer, Non-small cell lung cancer, Colorectal cancer, Endometrial cancer, Transcriptional misregulation in cancer, Huntington disease, Parkinson disease, Alzheimer disease, Non-alcoholic fatty liver disease (NAFLD), Infectious disease, Chagas disease (American trypanosomiasis), Signaling by FGFR1 in disease, SMAD4 MH2 Domain Mutants in Cancer. Modules without functional enrichment were excluded.

### Novelty scores of modules

Intuitively, the novelty score of a tumour module corresponds to how much of the module is found in normal modules.

First, we defined the raw novelty of a tumour module *A* as the minimum number (*N*) of normal modules that share 50% of the genes of the tumour module *A*. To calculate this, normal modules derived from the matched normal tissue are sorted by decreasing order of the number of genes in common with the tumour module. Then, starting from the module with the largest overlap, we count the number of normal modules that would be needed to achieve the 50% cut-off. This number represents the raw novelty score. The higher the novelty score, the greater number of normal modules were needed to achieve the cut-off, and therefore the tumour module is more specific to tumour samples.

To normalize for the association between novelty and the number of genes in the tumour module, we calculated the ratio between the raw novelty score of a module and the module size (*S*) (Equation 1).

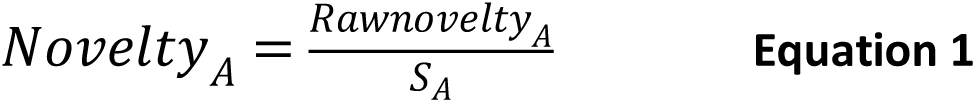

Modules were subsequently classified as being of High, Moderate or Low novelty by using the 1/3 and 2/3 quantiles as cut-off. As a result, “High novelty” modules are the most tumour-specific, and “Low novelty” modules are modules that are largely also found in normal samples.

### Level of expression of modules

The level of expression of modules was determined using ssGSEA^43^ from the GSVA package^34^ where genes of each sample are ranked based on their levels of expression. Higher scores are given when the genes of a module are consistently highly ranked.

### Somatic mutation analysis

A list of cancer drivers was obtained from the COSMIC Cancer Census database (Version Dec 21 23_46_30 2022)^44^ Genes were classified based on the disease where they have been reported to the drivers as per COSMIC, and were used in the analyses of the corresponding tumour type. Modules without cancer census genes were excluded from the analysis.

Point mutation and copy-number data for the tumour samples were obtained from TCGA and filtered as we have done previously^24^. A gene was considered to be recurrently point mutated if it had a missense or LoF mutation in at least 3 patients, and a non-synomymous to synonymous ratio less than 1. A gene was considered recurrently amplified or deleted if this occurred in at least 10% of the samples in the individual tumour cohort. Only focal amplifications and deletions were considered. We classified a gene as being focally amplified or deleted genes if their copy-number segment was in the upper quantile (0.25) of the distribution of the mean fraction of copy-number aberrant chromosomes across patients for each chromosome.

### Gene centrality

To calculate gene centrality, we first calculated the degree of genes in the co-expression networks, which was calculated as the sum of weights of the gene with all other genes in the network. To account for differences in the number of genes and overall strength of modules, the degree values were then normalized by dividing by the median degree of the module. To ensure the values were comparable across tumour types, genes were ranked by their normalized degree values. Subsequently, the ranks were divided by the size of the module, resulting in ‘relative’ ranks, with the maximum value being 1. A value of 1 was subtracted from the resulting values. Higher values would represent genes with a higher centrality, and lower values those that were more peripheral.

### Prostate cancer dataset

The RNAseq prostate adenocarcinoma dataset obtained from TCGA as described above. Tumour samples were further classified based on pathological annotations. Samples belonging to Grade Groups 1-3 (Gleason patterns 2+4, 3+3, 3+4, 4+3) where deemed to be “Low grade” and those of Grade Groups 4-5 (Gleason patterns 4+4, 3+5, 5+3, 5+4, 4+5, 5+5) where classified as “High grade”. This resulted in 45 low grade samples and 7 high grade samples, each with a matched normal sample.

Gene co-expression modules were derived from each sample set independently. Novelty scores were calculated comparing modules from low grade samples and those from normal samples, as well as from high grade samples to low grade samples. Modules were classified as high, medium or low novelty as described above using both sets of modules.

### Benign skin nevi and melanoma dataset

Gene expression data was obtained from Badal et al^41^ and downloaded from https://www.ncbi.nlm.nih.gov/geo/query/acc.cgi?acc=GSE98394. This dataset included 27 nevi samples and 51 primary tumour samples, however, one tumour sample was excluded during pre-processing as it was an outlier. Gene co-expression modules were derived from each sample set independently. Novelty scores were calculated comparing modules from primary and nevi samples. Modules were classified as high, medium or low novelty as described above.

### Benign and malignant pheochromocytomas

Gene expression data was obtained from previously published work^42^. This dataset included 17 normal samples from healthy adrenal tissue, as well benign and malignant samples from three subtypes of pheochromocytomas: MAML (9 benign and 8 malignant samples), SDH (44 benign and 36 malignant samples), and VHL (77 benign and 8 malignant samples).

Gene co-expression modules were derived from each sample set independently. Novelty scores were calculated comparing modules from benign and normal samples, as well as malignant and benign samples. Modules were classified as high, medium or low novelty as described above.

### Random Forest models

Single-sample GSEA scores (ssGSEA) for the 23 WGCNA modules derived from the TCGA glioblastoma (GBM) cohort were calculated for each RSEM-normalized RNA-sequencing sample from the TCGA GBM and low-grade glioma (LGG) cohorts, using the GSVA package as described above. Random Forest models were trained on ssGSEA scores and tested for their accuracy in classification of LGG and GBM samples. A 70/30 split was used for training and testing, repeated for 100 random splits of samples for training and testing. Area Under the Curve (AUC) for the Receiver Operating Characteristic (ROC) was recorded to assess model accuracy and mean decrease in Gini coefficients for each module were recorded to measure variable importance. For validation, the mRNAseq 693 (batch 1) RSEM RNA-sequencing gene expression dataset was downloaded from The Chinese Glioma Genome Atlas (CGGA) (http://www.cgga.org.cn/index.jsp)^45^. The dataset contained 693 samples of both primary and recurrent low-grade gliomas and glioblastomas. Random Forest model training and testing for accuracy in classification of LGG and GBM samples was conducted as described for TCGA data. Model training and performance assessment was performed in R 3.6.3 using the randomForest (v4.6-14) and pROC (v1.17.0.1) packages.

## Acknowledgements

DLG was supported by a Victorian Cancer Agency fellowship (MCRF17005) and the Peter MacCallum Cancer Foundation. AST was supported by National Health and Medical Research Council Ideas Grants 2003115 and 2003887. AST is a Prostate Cancer Foundation Young Investigator. We thank the Research Computing Facility at the Peter MacCallum Cancer Centre for providing the infrastructure and support to carry out this project. We thank Dr. Kimberly Bussey, Prof. Kieran Harvey and Prof. David Bowtell for critical reading of the manuscript. We thank Prof. Mark Rosenthal for inspiration for the brain cancer analysis and Dr. Benjamin Goudey and Dr. Thomas Conway for suggesting the random forest approach. We thank Mikhail Dias for designing diagrams. We thank Anna Small and Jodie Kirkland for assistance with manuscript preparation.

## Code availability statement

Code to reproduce all analysis is available at https://github.com/cancer-evolution/Evolutionary_analysis_of_coexpression_modules

