## Supplementary material for "Disruption of metazoan gene regulatory networks in cancer alters the balance of co-expression between genes of unicellular and multicellular origins"

### Supplementary Figures

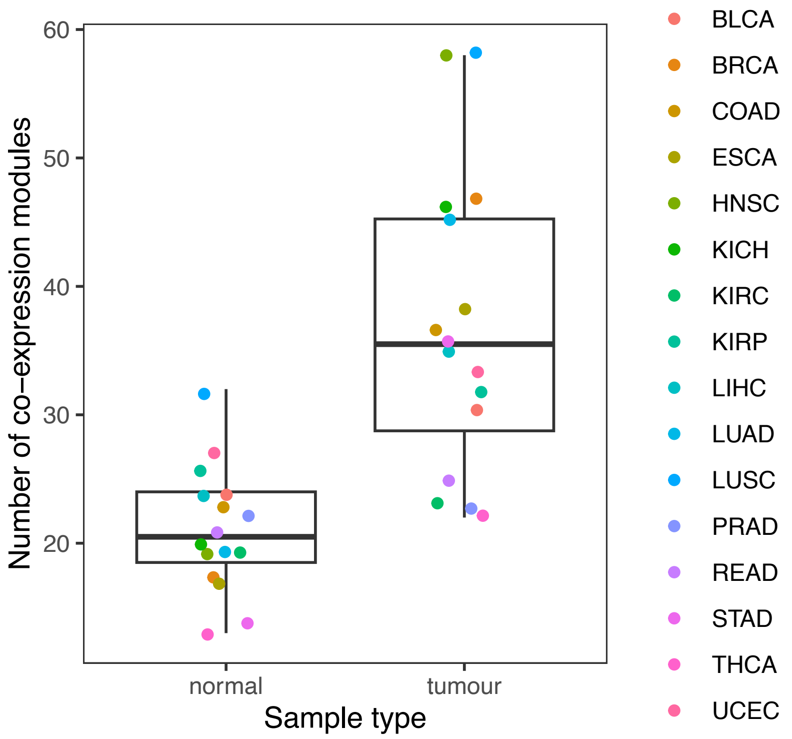


**Figure S1. Number of co-expression modules for cohorts with only matched tumour and normal samples.**

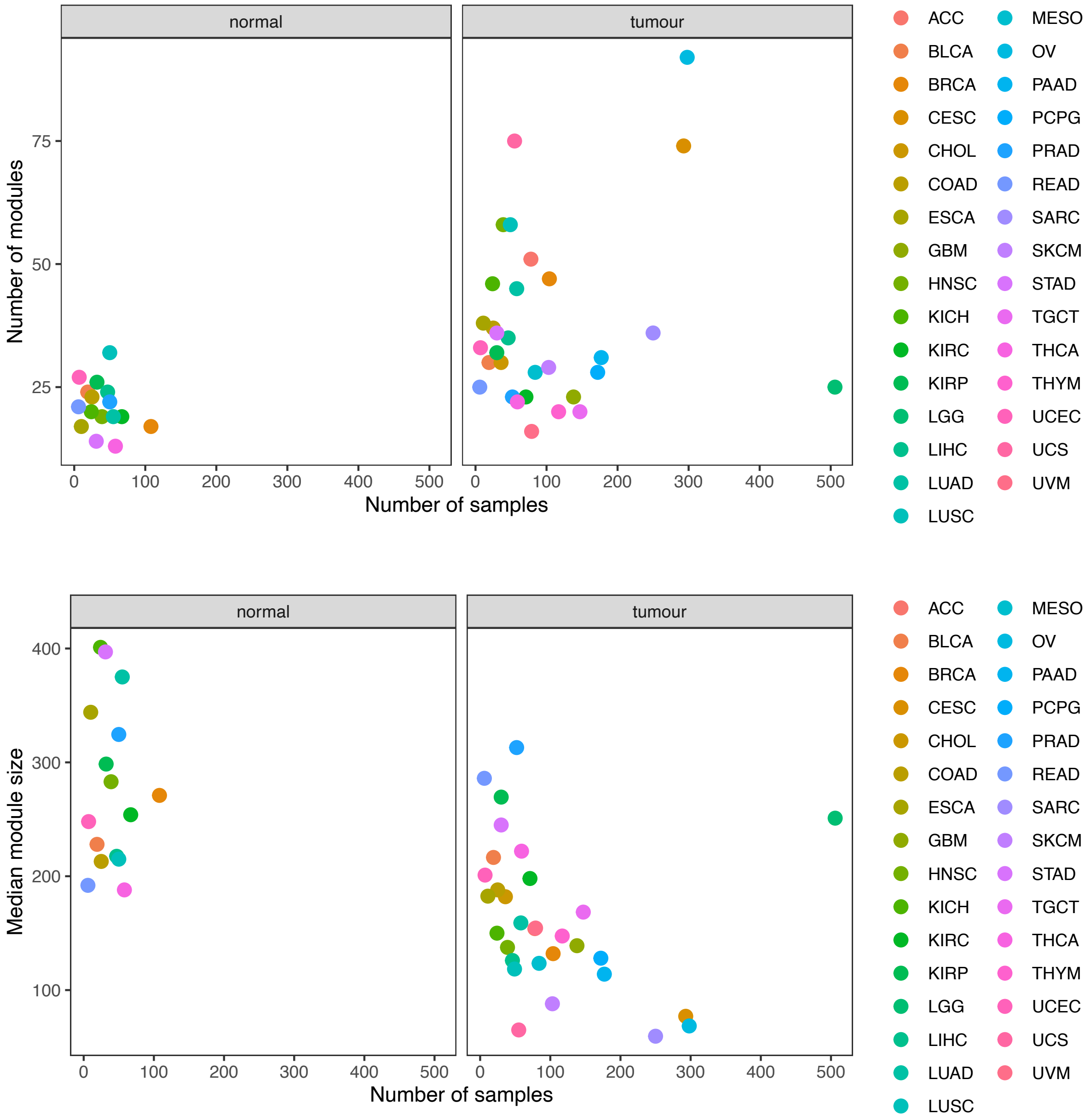


**Figure S2. Correlation between number of modules/modules sizes and cohort size.** Spearman correlation values of top panel: Normal= -0.32, Tumour = -0.12. Spearman correlation values of bottom panel: Normal = -0.0059, Tumour = -0.53.

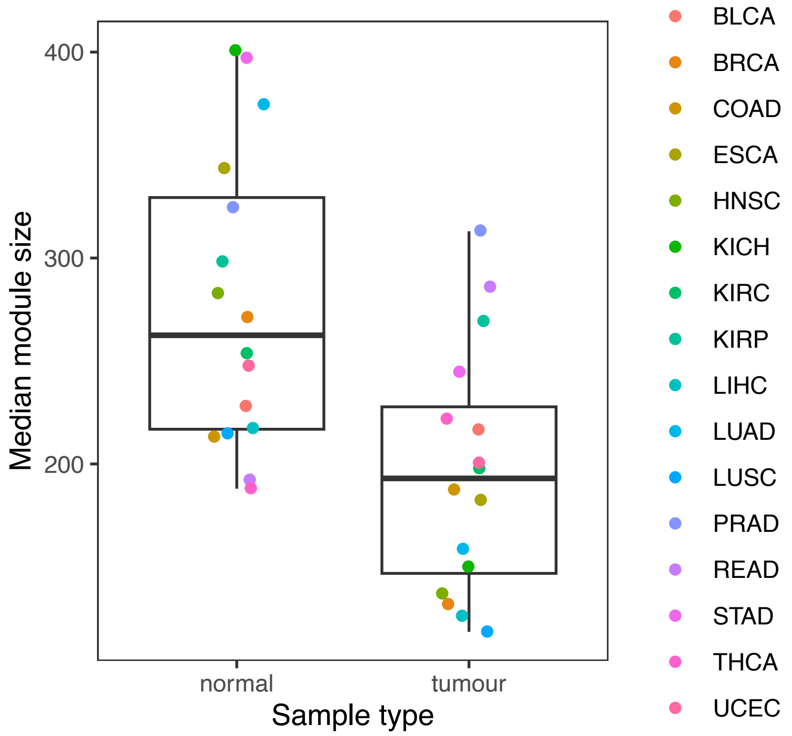


**Figure S3. Module sizes for cohorts with only matched tumour and normal samples**

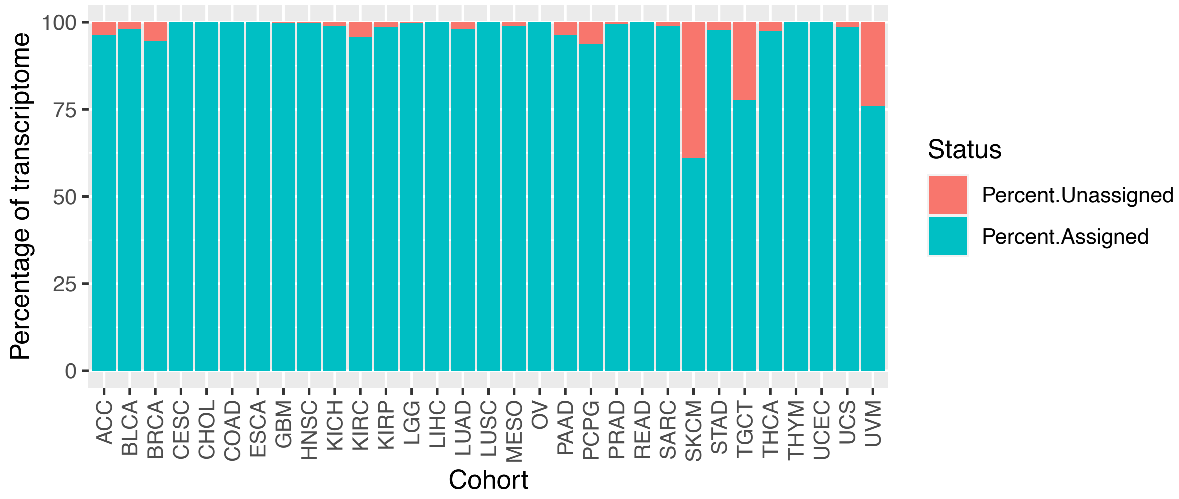


**Figure S4. Percentage of genes assigned to modules per cohort.** The percentage of unassigned genes correspond to those in the 'grey' module.

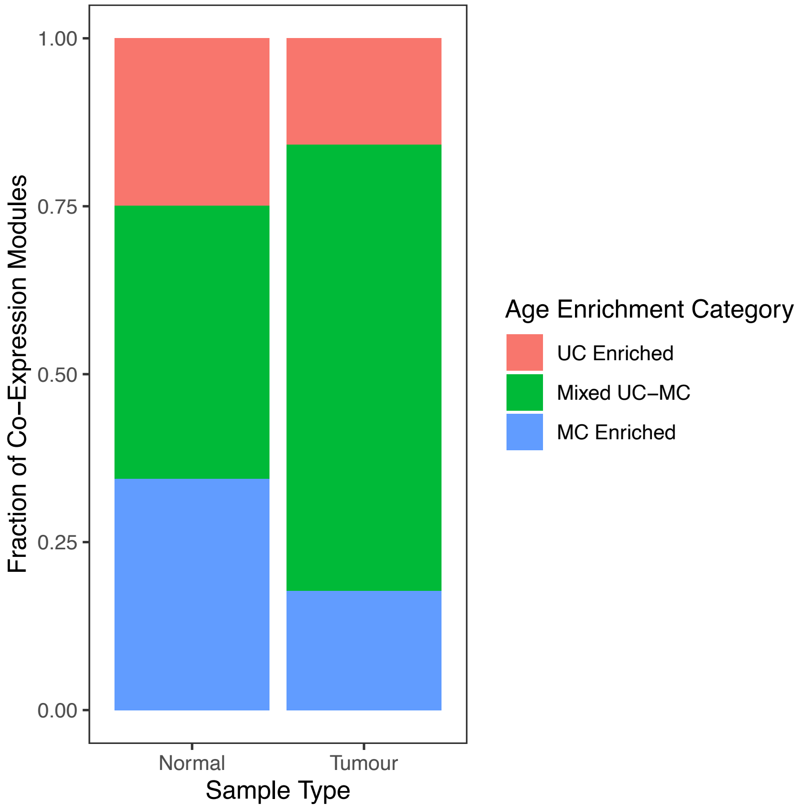


**Figure S5. Fraction of age enrichment categories among tumour and normal modules**

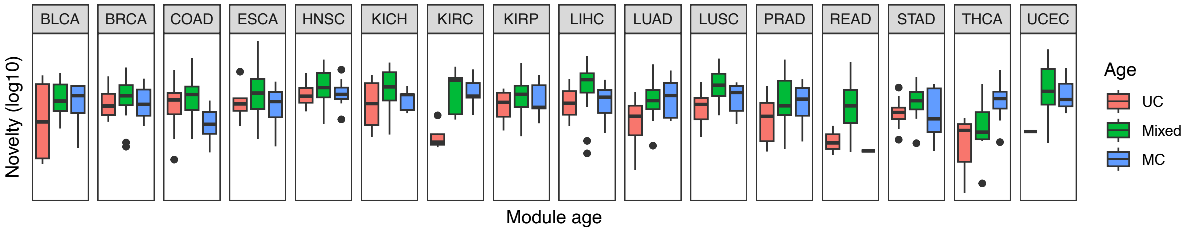


**Figure S6. Differences in Novelty scores for UC-enriched, Mixed UC-MC and MC-enriched tumours across all cohorts**

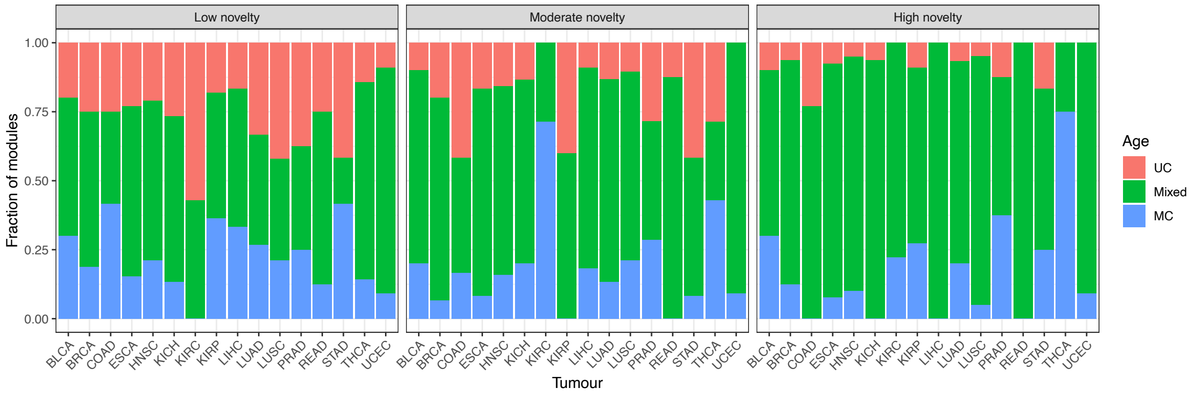


**Figure S7. Fraction of modules of each age per novelty category in each tumour cohort.**

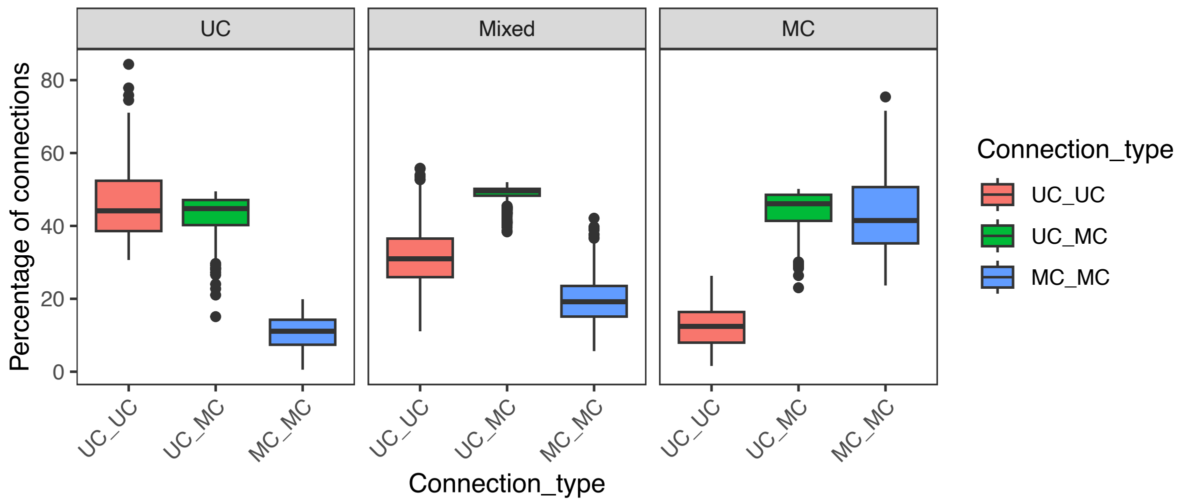


**Figure S8. Percentage of connections that are UC-MC in Mixed UC-MC, UC-enriched and MC-enriched modules.**

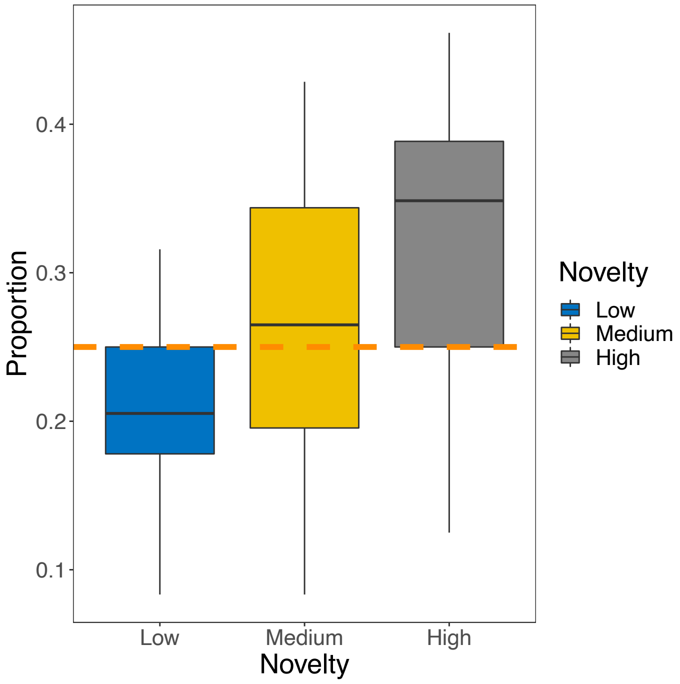


**Figure S9. Proportions of high, medium and low Novelty modules with ssGSEA scores in the upper quartiles of all ssGSEA scores calculated for their respective tumour cohorts.**

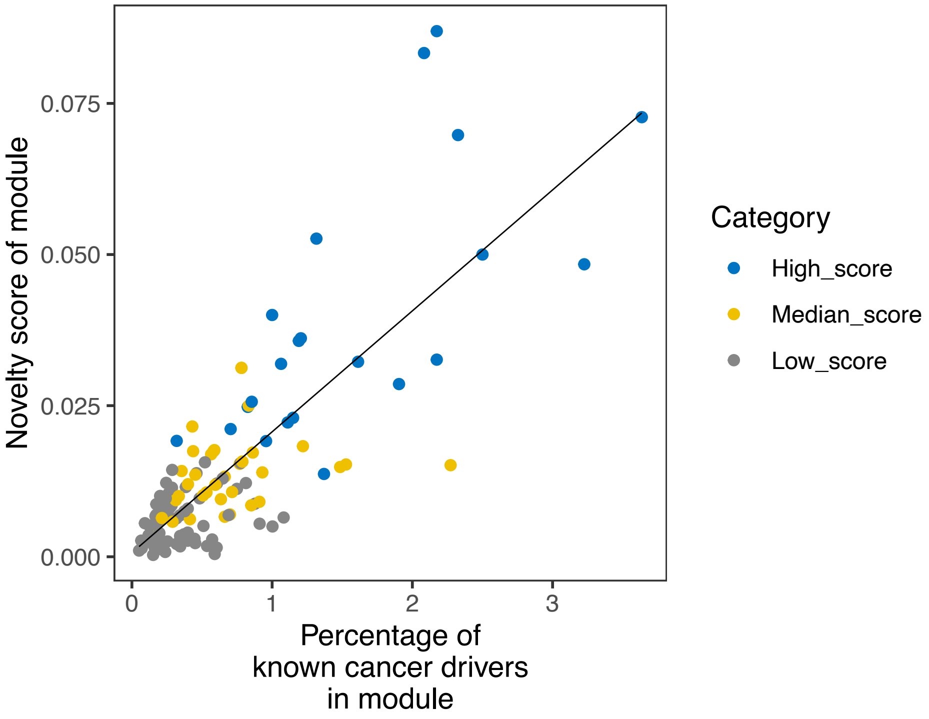


**Figure S10. Correlation between module novelty score and percentage of known cancer driver genes from COSMIC Cancer Census gene list, stratified by module novelty.**

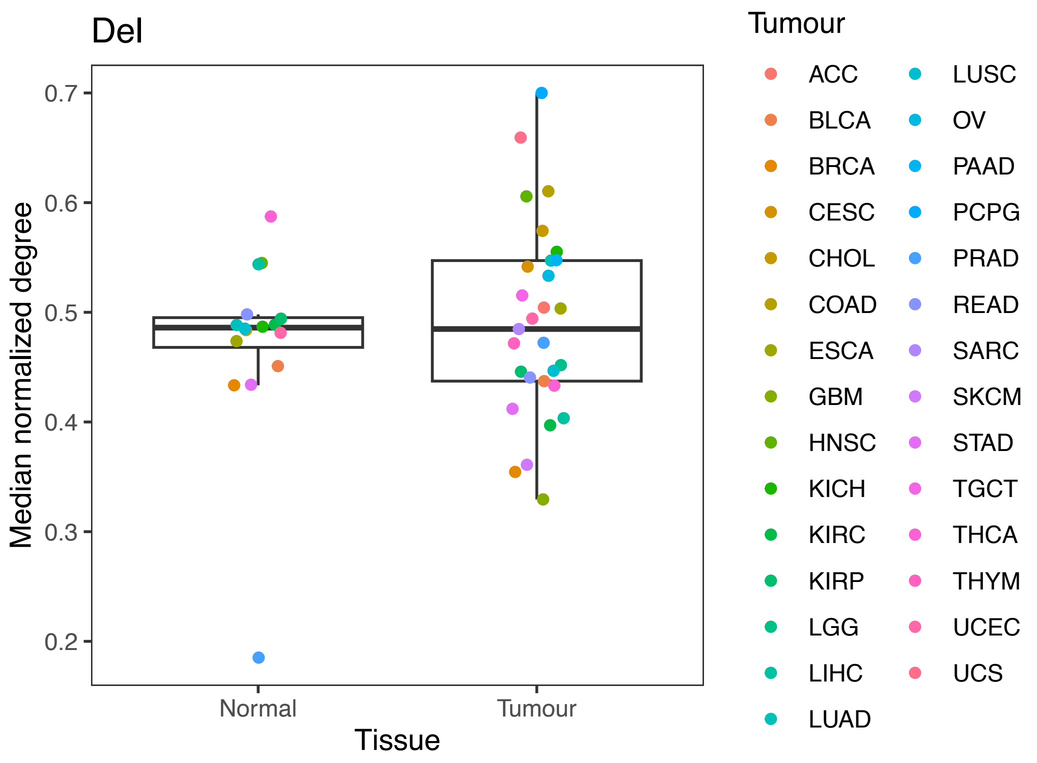


**Figure S11. Normalized degree of recurrently amplified genes within WGCNA co-expression modules from normal (left) and tumour (right) cohorts, coloured according to tumour type.**

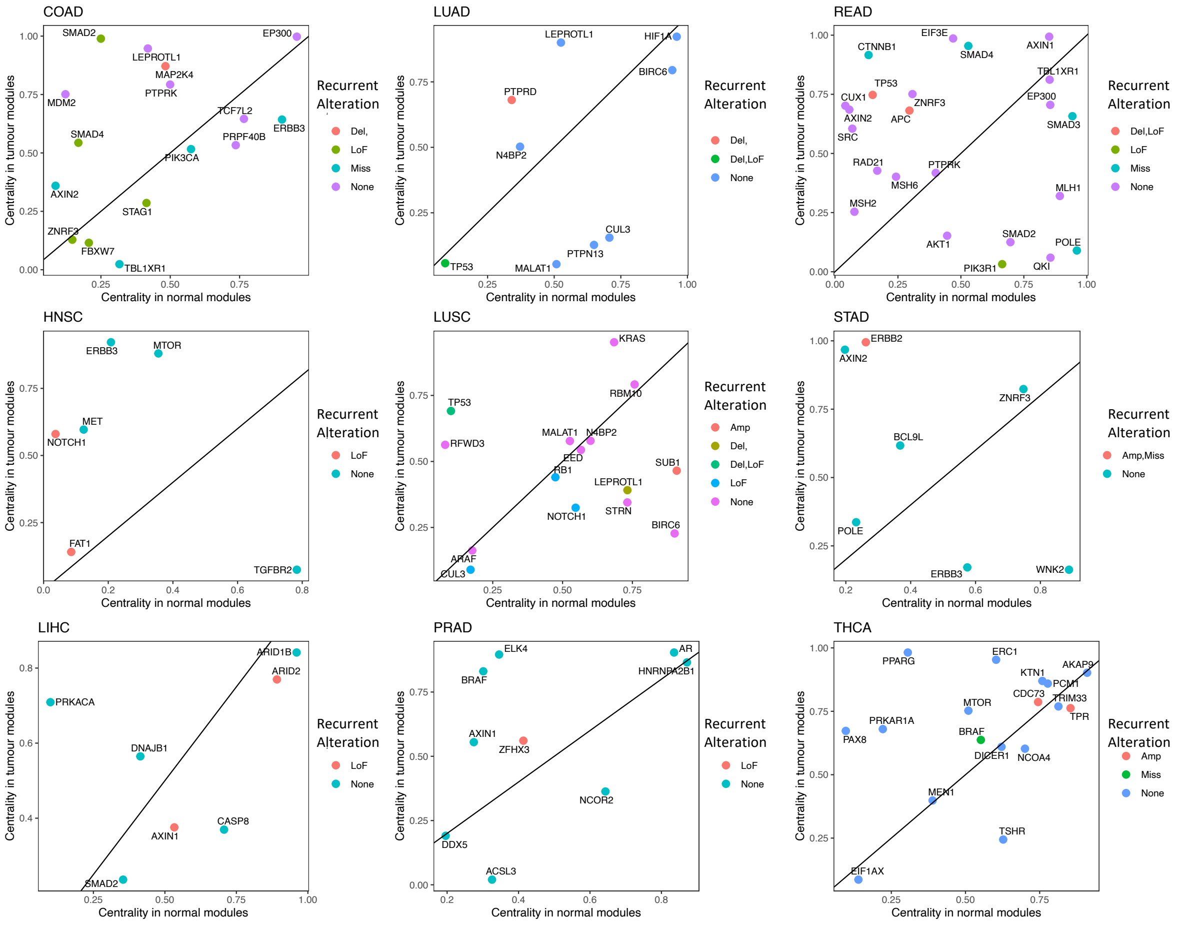

**Figure S12. Changes in centrality for driver genes in other solid tumour cohorts from TGCA**

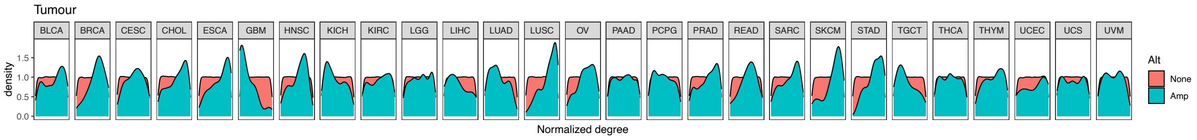


**Figure S13. Centrality of recurrently amplified and non-mutated genes in modules from tumour cohorts**

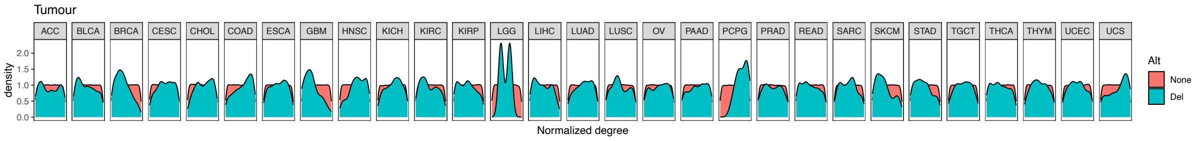


**Figure S14. Centrality of recurrently deleted and non-mutated genes in modules from tumour cohorts**

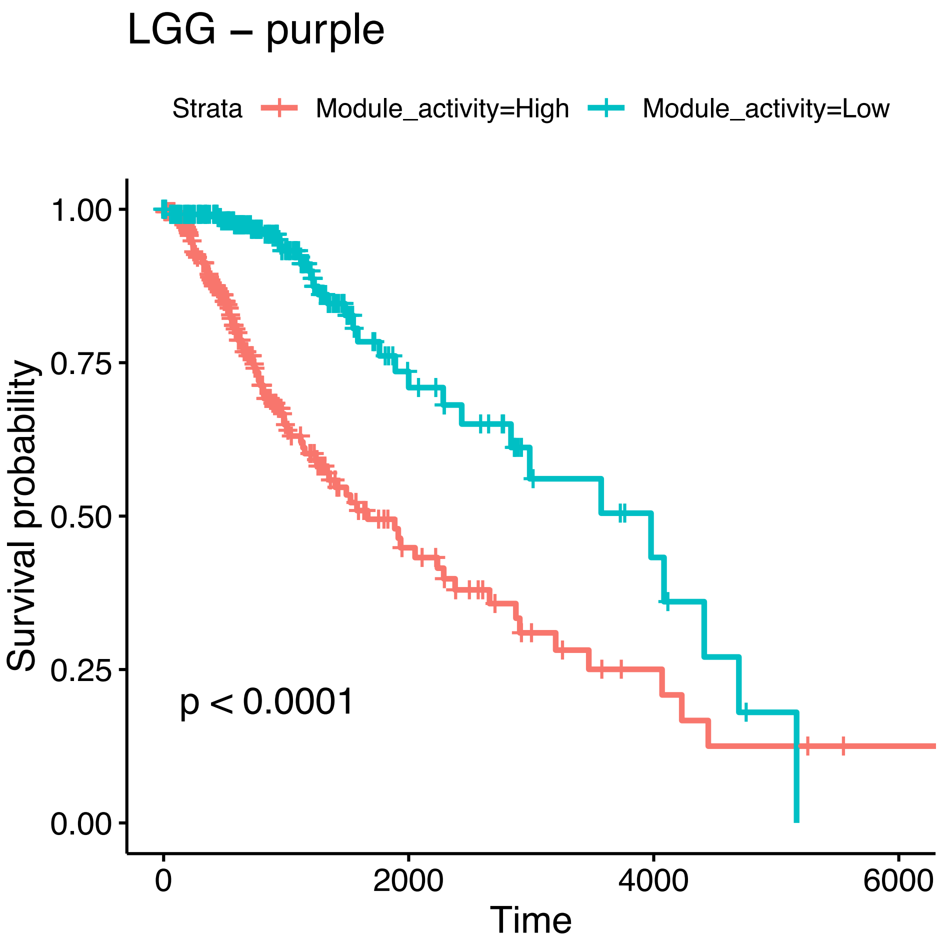


**Figure S15. Survival curves for low vs high expression of LGG purple module.** Patients were classified as either having high (equal or above the median) or low (below the median) activity of the Purple module. N=514 patients.

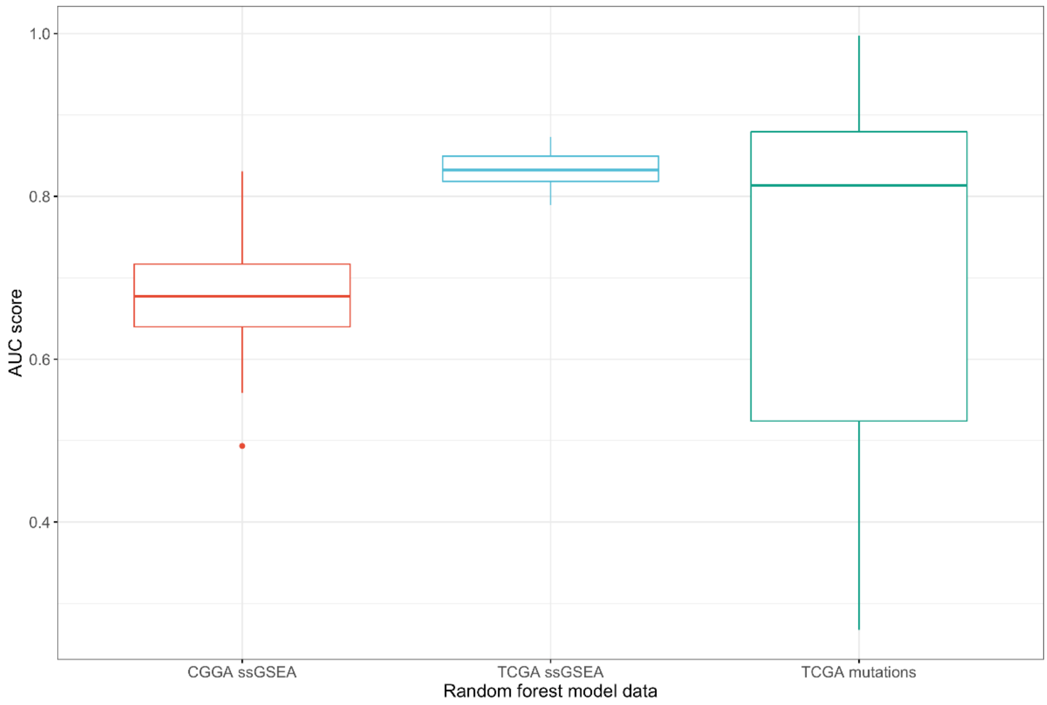


**Figure S16.** **AUCs of Random Forest models trained on mutation status from TCGA (green), ssGSEA scores of GBM modules tested on CGGA (red) and TCGA (blue) datasets.**

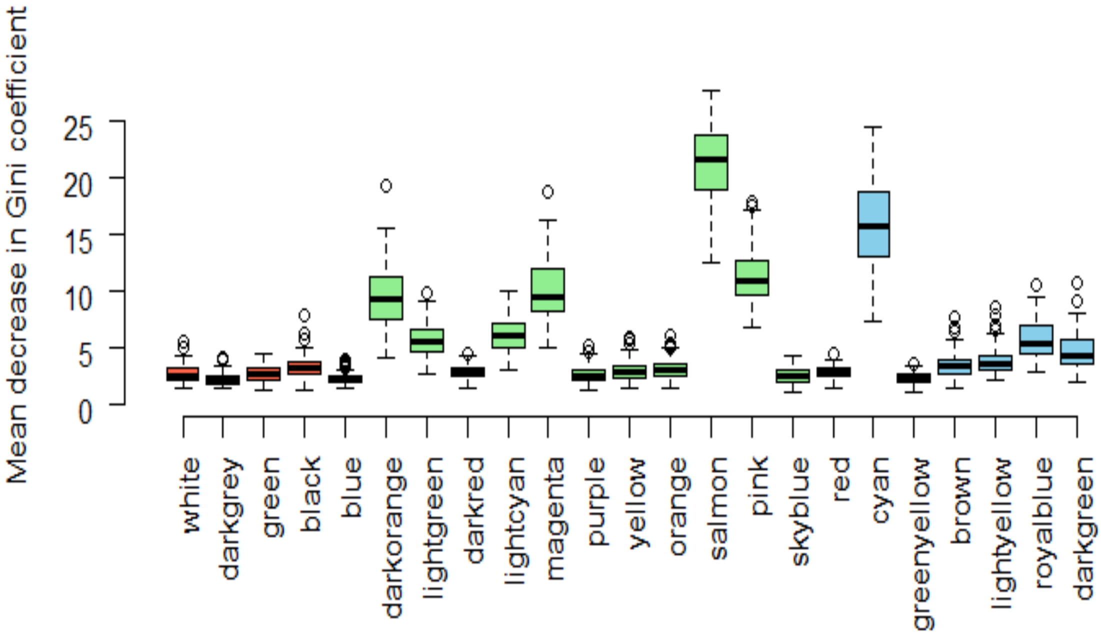


**Figure S17. Mean decrease in Gini Coefficient for GBM modules from TCGA from 100 iterations of Random Forest models tested on RNA-seq data from the CGGA.**

### Supplementary Tables

**Table S1. Number of modules for Tumours & Normal cohorts by age enrichment category**

| | | Number of Co-expression Modules | | | | | | |
|---|---|---|---|---|---|---|---|---|
| **Module Age Enrichment Category** | **Sample Type** | **Minimum** | **1st Quartile** | **Median** | **Mean** | **3rd Quartile** | **Maximum** | **Total** |
| Unicellular-enriched | Tumour | 1 | 4 | 6 | 5.97 | 8 | 12 | 185 |
| | Normal | 1 | 3.75 | 5.5 | 5.25 | 7 | 8 | 84 |
| Mixed Unicellular-Multicellular | Tumour | 6 | 14.5 | 20 | 24.97 | 31 | 82 | 774 |
| | Normal | 2 | 6 | 9 | 8.56 | 11 | 16 | 137 |
| Multicellular-enriched | Tumour | 1 | 6 | 7 | 6.68 | 8 | 10 | 207 |
| | Normal | 2 | 6.75 | 7 | 7.25 | 8 | 13 | 116 |

### Supplementary Data

- Phylostrata assignments for all human genes
